# Parasitology, Poverty and Prevention: is there any relationship between the three P? Is it possible to eradicate Parasitic diseases without eliminating Poverty?

**DOI:** 10.1101/544007

**Authors:** Guyguy Kabundi Tshima, Paul Madishala Mulumba

## Abstract

**Context:** Talking about Poverty is not obvious without examples, I would like to understand the link between Parasitology, Poverty and Prevention (the three P). I explain the three P by saying that there is four level of knowledge in Parasitology and the fourth level is the integration with other disciplines including virology with preventive measures, nutrition aspects with denutrition leading by some parasites as Ascaris, economy involving patient’s income and Poverty. As a reminder, the first level in Parasitology is the knowledge of the parasitic cycle with an emphasis on the mode of contamination, the second level is that of the implementation of technical or diagnostic means to identify the parasite in the laboratory or the bench and the third level is that of treating infected cases diagnosed in the laboratory.

**Objective:** The objective of this work is to contribute to reach the first sustainable development goal i.e. no Poverty. Specifically, this manuscript aimed to evaluate poverty with the protective measures against the harmful effects of mosquitoes that contribute to the quality of care given to patients of the University Hospital of Kinshasa (UHK).

**Findings:** Residual mosquito capture, carried out in 31 randomly selected rooms per block and per level in hospital departments, presented the number of 1,144 female mosquitoes (845 *Culex*, 207 *Anopheles* and 62 Aedes). Overall considered, the Mean Mosquito Density (MMD) was 36.2 / mosquito per room (6.9 *Anopheles* / room, 29.1 *Culex* / room and 2.1 *Aedes* / room with an extreme between 0 and 144 mosquitoes / room. The lowest MMD (6.2 mosquitoes / room) was observed in Block II (clinical biology and microbiology laboratories, delivery and private hospitalization rooms) compared to other hospital blocks that had the highest MMD and statistically identical (ranging between 29.2 and 45.5 mosquitoes / room).

Our observations give a good idea of Poverty inside this hospital and where to concentrate in the prevention of malaria transmission within the hospital. Regardless of the block considered, it was the ground floor with an MMD of 52.8 mosquitoes / room which were the most dangerous places compared with the first and second floors with MMD respectively 17.6 and 25.6 mosquitoes / room.

**Conclusion:** In conclusion, the insufficiency of the UHK anti-mosquito measures was obvious. These should be applied without delay to prevent the risk of infection transmission by mosquitoes, even within the hospital, of hepatitis B virus and strains of *Plasmodium falciparum*, sometimes highly virulent, which may be concentrated there.

**Limits:** We were on the right track and this study needs more research because of its limitations: we investigate and did not find if any of the mosquitoes collected were infected; we did not investigate if the hospital had any patients with a mosquito transmitted disease in the rooms where the mosquitoes were collected.

**Recommendation:** The recommendation is if it is not possible to eradicate parasitic diseases as malaria without eliminating poverty, then we need to eliminate them both.

## Résumé

**Contexte:** Parler de la pauvreté n’est pas évident sans exemples, je voudrais comprendre le lien entre parasitologie, pauvreté et prévention (les trois P). J’explique les trois P en disant qu’il existe quatre niveaux de connaissance en parasitologie et que le quatrième niveau est l’intégration à d’autres disciplines, notamment la virologie avec des mesures préventives, les aspects nutritionnels avec la dénutrition observée avec certains parasites comme l’Ascaris, l’économie impliquant le revenu des patients et la pauvreté. Pour rappel, le premier niveau en parasitologie est la connaissance du cycle parasitaire en mettant l’accent sur le mode de contamination, le second niveau est celui de la mise en œuvre de moyens techniques ou diagnostiques permettant d’identifier le parasite en laboratoire ou sur le banc et le troisième niveau est celui du traitement des cas infectés diagnostiqués en laboratoire.

**Objectif:** L’objectif de ce travail est de contribuer à la réalisation du premier objectif de développement durable, c’est-à-dire l’absence de pauvreté. Plus précisément, ce manuscrit visait à évaluer la pauvreté grâce à des mesures de protection contre les effets nocifs des moustiques qui contribuent à la qualité des soins prodigués aux patients des Cliniques Universitaires de Kinshasa (CUK).

**Résultats:** La capture résiduelle de moustiques, effectuée dans 31 locaux choisis au hasard par bloc et par niveau dans les services hospitaliers, a présenté le nombre de 1 144 moustiques femelles (845 *Culex*, 207 *Anopheles* et 62 *Aedes*). Dans l’ensemble, la densité moyenne des moustiques (DMM) était de 36,2 / moustiques par local (6,9 *Anopheles* / local, 29,1 *Culex* / local et 2,1 *Aedes* / local avec une extrême entre 0 et 144 moustiques / local) ont été observés dans le bloc II (laboratoires de biologie clinique et de microbiologie, salles d’accouchement et d’hospitalisation privée) par rapport aux autres blocs hospitaliers qui avaient la plus haute DMM statistiquement identiques (allant de 29,2 à 45,5 moustiques / local).

Nos observations donnent une bonne idée de la pauvreté à l’intérieur de cet hôpital et du lieu où se concentrer dans la prévention contre le paludisme. Quel que soit le bloc considéré, c’étaient les rez-de-chaussée avec une DMM de 52,8 moustiques / local qui étaient les endroits les plus dangereux par rapport aux premiers et deuxièmes étages avec DMM respectivement 17,6 et 25,6 moustiques / local.

**Conclusion:** En conclusion, l’insuffisance des mesures anti-moustiques des CUK était évidente. Celles-ci doivent être appliquées sans délai pour prévenir le risque de transmission par les moustiques, même à l’intérieur de l’hôpital, du virus de l’hépatite B et de souches de *Plasmodium falciparum*, parfois extrêmement virulentes, qui peuvent s’y concentrer.

**Limites:** Nous étions sur la bonne voie avec la notion de risque qui n’était pas confirmé donc cette étude qui nécessite davantage de recherche en raison de ses limites: nous avons enquêté et n’avons trouvé aucun des moustiques collectés infectés; aussi nous n’avons pas cherchéà savoir si l’hôpital avait des patients atteints d’une maladie transmissible par les moustiques dans les salles où les moustiques avaient été collectés.

**Recommandation:** La recommandation est la suivante: s’il n’est pas possible d’éradiquer les maladies parasitaires comme le paludisme sans éliminer la pauvreté, nous devons éliminer tous les deux.

**Mots-clés:** nuisance culicidienne, Cliniques Universitaires de Kinshasa, République Démocratique du Congo.

## Introduction

At the UN Summit of September 2015, the Member States of the United Nations adopted the 2030 Agenda for Sustainable Development with a set of Sustainable Development Goals (SDGs) that **aim to end extreme poverty, promote equity and opportunity for all, and protect the planet** (https://www.un.org/sustainabledevelopment/development-agenda/). SDGs are universal goals accepted by all countries and applicable to all – both rich and poor, taking into account different national realities, capacities and levels of development and respecting national policies and priorities (https://www.connectyet.org/sdg/category/about-sdg) worldwide and the Democratic Republic of the Congo cannot miss this opportunity to open itself to development also the University Hospital of Kinshasa (UHK), located on the hill of Mount Amba, in the southern suburbs of the city of Kinshasa, that occupies the top of the health system of the Democratic Republic of the Congo (DRC). Compared to the rest of the city, hospital environments are a real reactor that concentrates most pathogens at a very high-density level. As a result, the biosecurity problem is acute. Conventional prophylactic measures to prevent nosocomial infections do not usually include those intended for protection against mosquito’s nuisance, when they should. This work set itself the objective of assessing the level of mosquito’s nuisance that reigns in this important care institution in the Democratic Republic of the Congo.

## Materials and methods

### Inclusion and exclusion criteria

To be eligible, a block should be accessible to the survey in all its parts. At each level of the selected block, to be eligible, a room should be accessible to the investigators. The UHK have 4 hospital blocks, 1 consultation block (block C) and 1 technical block. In its hospital part, each block has 4 levels: 1 basement, 1 ground floor and 2 floors. Block C has only 2 levels. Three of the four hospital blocks (blocks I, II and IV) and the block of consultations (block C) met this condition. Block III, which has at the last level a hospitalization service, was not retained because of the ineligibility of these lower levels which contain sensitive services: radiology and nuclear medicine on the ground floor, the resuscitation and operating rooms on the 1st floor. Also excluded: bedridden patients’ rooms, operating rooms, delivery rooms, laboratories, imaging department, intensive care units, stores, basements and pharmacy.

### Sampling

Showing the documents authorized by the UHK authorities, we proceeded with the identification of the places and the census of the premises in which the mosquitoes capture was to take place. Out of a total of 343 local respondents considered eligible for our survey, one in ten rooms was selected using the random number table by level and block. Some additional premises were drawn in anticipation of the possible replacement of premises that would have lost their eligibility in the meantime.

### Mosquito capture

The capture of mosquitoes, carried out in 6 sessions, took place in 31 premises instead of 34 expected. We used insecticide bombs charged permethrin spray, a synthetic pyrethroid. Before spraying insecticide into some rooms, the following precautions were taken: the occupants were asked to leave, taking away their valuables, all the windows were closed, dishes, food and drinks, were evacuated. Furniture and tiles were covered with white sheets.

Masked and gloved, we started spraying from the outside to the inside, first to the open door that we close behind us while progressing on the bottom of the room to spray every possible corner. Spraying a room, depending on its size, lasted from 20 to 40 seconds. We left the room closing the door behind us. After waiting 10 minutes, the door and windows were opened again for 5 minutes to ventilate the treated room. Then, using pliers, we proceeded to collect mosquitoes fallen on the sheets. Those fallen under the furniture were searched with a flashlight.

### Entomological examination

Captured mosquitoes were brought back to the laboratory for identification. After exclusion of males, females were sorted and enumerated by genus.

### Statistical analyzes

The comparison of mean mosquitoes’ densities (MMD) by local was done by means of the analysis of variance (Snedecor test F) following a general linear model in which the blocks, the levels played the role of fixed factors, and, no random draw, while the premises were taken as a random factor because drawn.

Given the large variability of the data (mean-proportional dispersion), this analysis was performed on the logarithms of the data to which “1” was added to account for zeros. In this comparison of MMD, *Anopheles*, a vector of malaria, was opposed to other *Culex* and *Aedes* taken together, given their small size.

The significance of the results was assessed at the 5% probability level of significance.

## Results

### Comparison of the nuisance between the blocks

The collection, carried out in 31 rooms, brought back 1,144 female mosquitoes including 207 *Anopheles*, 845 *Culex* and 62 *Aedes*, respectively 19.2; 74.9 and 5.9% of the workforce. Overall, MMD per local was 6.9 for *Anopheles* and 29.1 for *Culex* and 2.1 for Aedes (Table 1).

The comparison of the MMD between the blocks did not show a significant difference (p = 0.327). However, orthogonal contrast analysis showed that Block II had the lowest MMD compared to the others (p = 0.001).

**Table 1.** MMD study by block and by type (arithmetic mean ± 1 standard deviation)

| BLOCK | GENUS |  | TOTAL |
| --- | --- | --- | --- |
|  | *Anopheles* | *Culex+Aedes* |  |
| I | 7.7±15.1 (13) | 37.8±38,1 (13) | 45.5±64.4 (13) |
| II | 2.0±1.8(4) | 4.3±2.2 (4) | 6.2±4.4 (4) |
| VI | 7.7±5.5 (10) | 32.8±31.7 (10) | 40.4±51.2 (10) |
| C | 9.5±5.7 (4) | 19.8±14.9 (4) | 29.2±23.6 (4) |
| **TOTAL** | **7.1±10.3 (31)** | **29.1±31.9 (31)** | **36.9±52.0 (31)** |

- 355 • Block I: Ground floor (Pediatrics), 1st floor (Maternity, Neonatology), 2nd floor (child surgery, specialties and gynecological hospitalization) - 357 • Block II: Ground floor (clinical biology, microbiology); 1st floor (delivery rooms, gynecology operating rooms); 2nd floor (private hospitalization). - 359 • Block IV: Ground Floor (Hospitalization Internal Medicine, Intensive Care); 1st stage (hospitalization surgery, hospitalization internal medicine); 2nd floor (hospitalization surgery, hospitalization internal medicine).

**Table 2.** Study of MMD between blocks (orthogonal contrast analysis)

| CONTRAST | PROBABILITY |
| --- | --- |
| (Block II) vs (Others) | 0.001 |
| (Block I) vs (Blocks IV and C) | 0.457 |
| (Block IV) vs (Block C) | 0.854 |

### Comparison of mosquito’s density by level

Overall, there was no significant difference between levels (p = 0.437). However, orthogonal contrast analysis showed that the ground floors had a significantly higher MMD than those observed in the floors (p = 0.047). Between floors, the difference was not significant (p = 0.091); (Table 4) The ground floor with an average of 52.8 mosquitoes / local, while the first and second level had respective values: 17.6 and 25.6 mosquitoes / local. The interaction analysis showed that the difference in MMD between the levels was the same from one block to another (p = 0.425).

### Comparison of MMD by local

Overall, there was no significant difference in MMD between premises (p = 0.926). However, the study of interactions showed the difference between premises was not the same, nor from one block to another (p <0.0001) nor from one level to another (p = 0.0001). The difference observed between the species was identical whatever the place (p = 0.712), but not from one block to another (p = 0.001) (Table 3).

**Table 3.** MMD study according to the levels (arithmetic average ± 1 standard deviation), (the number of mosquitoes is in parentheses)

|  | CULICIDIAN GENUS |  |  |
| --- | --- | --- | --- |
| LEVEL | *Anopheles* | *Culex+Aedes* | **TOTAL** |
| Ground floor | 10.1±14.0 (16) | 42.7±39.6 (16) | 52.8±67.2 (16) |
| 1st floor | 3.2±3.6 (9) | 14.3±14.3 (9) | 17.6±24.0 (9) |
| 2<sup>nd</sup> floor | 6.0±4,1 (6) | 19.7±15.5 (6) | 25.6±26.0 (6) |
| **TOTAL** | **7.1±10.3 (31)** | **29.1±31.9 (31)** | **36.2±52.0 (31)** |

**Table 4.** Comparison of MMD between Blocks (Orthogonal Contrast Analysis)

| CONTRAST | PROBABILITY |
| --- | --- |
| Ground Floor vs (1<sup>st</sup> et 2<sup>nd</sup> floor) | 0.047 |
| 1<sup>st</sup> floor vs 2<sup>nd</sup> floor | 0.091 |

## Discussion

The work reports findings with implications for public health. Given the implication for patient care or public health, among many causes, I think poor countries deserve to be affected by parasitic diseases because of illiteracy, health negligence. It is observed that parasitology-virology diseases incidence is more common in poor regions of the world [1–5]. Any relationships in mechanisms of those diseases and poverty? On one hand malaria is often referred to as the epidemic of the poor whilst the disease is in large part determined mainly by climate and ecology [1], and not poverty, on the other hand, the impact of malaria takes its toll on the poorest [1], those least able to afford preventative measures and medical treatment and those living in suboptimal and poor hygiene and sanitation conditions that are indeed the root causes that define the ascariasis [2]. The risk of HIV transmission in theory is also involved in the poor settings [3] where when the international aid stops, disease prevalence become more prevalent as the case of trypanosomiasis [4]. In this context, MMD at CUK was 36.2 female / local with extremes mosquitoes ranging from 0 to 144 mosquitoes / local. This number represents potentially as many bites to which all people sharing the same premises were exposed.

If we consider, for example, an average of 10 people per room (patients and nurses included), each of them will be exposed to receive an average of 3.6 bites per person per night (PPN), or 5,114 PPN annually. This nuisance was mainly due to *Culex*, which alone accounted for 75.9% of the mosquito population caught. Compared with *Anopheles, Culex* was 4.1 times more numerous. The average number of PPN due to *Anopheles* would be 0.7 per local. This number should be increased to 5.4 in the extreme situations observed on the ground floor.

Thus, as evidenced by the results of this survey, the risk of malaria transmission within the UHK itself can not be neglected [5]. Of course, we investigated our capture, and we did not find any of the mosquitoes collected were infected; but ethically we had not the right to investigate if the hospital had any patients with a mosquito-transmitted disease in the rooms where the mosquitoes were collected. We know that the risk of transmission of parasitic infections (HIV, yellow fever, dengue fever, Japanese encephalitis, viral hepatitis, and filariasis), is very real for some patients, this study highlighted that the risk of infection remains for others purely theoretical [5].

As mosquitoes can transmit yellow fever, dengue fever and Japanese encephalitis, there is no evidence that they are capable of transmitting HIV. Indeed, it is established experimentally, by gene amplification techniques, that this virus disappears from the body of the mosquito after 1 to 2 days, time required for digestion of the blood meal. Consequently, since the virus can not survive the time required to invade the salivary glands and multiply due to the absence of CD4 receptors in arthropods, its cyclic transmission is therefore not possible [6]. There remains only the possibility of a mechanical transmission.

A mosquito that interrupted his meal on an HIV-positive patient, can he, when he immediately restarted on another subject, transmit the infection? Here again, it is established that HIV is inactivated within 20 minutes during its stay in the fallopian tubes. However, a mosquito that has interrupted its meal can not statistically restart it before 20 minutes [7]. Moreover, even if some viruses remain in the trunk, it has been calculated that for a mechanical transmission to be possible, the healthy subject must receive an inoculum containing at least 10 million virus particles. Due to a hypothetical inoculation of 10 copies of viral RNA per sting, it would take 1 million bites that can be expected to receive an individual during his stay, even extended to several years, UHK.

Regarding yellow fever, the mere presence of *Aedes aegypti*, which accounts for 5.6% of CUK’s culicidae fauna, is not enough to ensure the transmission of this disease. Indeed, the cycle of the yellow fever virus requires the interlocking of three cycles to pass the virus from the forest (monkeys) to the city (man). The biotope of Kinshasa, far away from the forests, is not conducive for the development of such a chain of transmission. On the other hand, the intrusion of a single case of yellow fever into the hospital is potentially dangerous for all the people who live there. Given the presence of the potential vector, draconian biosecurity measures must be taken without delay.

Regarding the transmission of hepatitis B (HBV) and (HBC) viruses by mosquitoes, there is a few studies that have focused on this issue. Regarding the transmission of HBV, the work of Blow et al [8] showed that this virus could be transmitted to humans through the droppings of mosquitoes, especially when they are forced to interrupt their meal on a subject infected and then resume immediately on another subject. In addition, the work of Zhang et al [9] on monkeys confirmed the possibility of cyclic transmission of HBV by mosquito saliva. Moreover, the work of Fouché et al [10], conducted on mosquitoes of the *Culex quinquefasciatus* species, demonstrated the existence of a vertical (transovarian) transmission of HBV infection from one generation to the next other.

Regarding the transmission of HCV, both mechanically and cyclically, the results of various studies remain divergent on this question. Indeed, there are those who support this idea, this is the case of Hassan et al [11] and Washenberger et al [12], while, for their part, Yeung et al [13] reject it. The risk of transmission of the Japanese encephalitis virus is theoretically possible even if no case has been reported so far. Indeed, the available vectors (*Aedes, Culex*) are legion in our environment and the UHK. As for the dengue, although the DRC is included in the distribution area of *Aedes aegypti*, dengue cases, which they are indigenous or imported, have never been reported to date. Dengue is a tropical pathology caused by 4 species of virus whose cycle involves humans and *Aedes aegypti*, an anthropophilic mosquito with diurnal activity. It must be admitted, however, that the hospitalization of a single case of dengue at UHK is justifiable for quarantine measures, especially for the haemorrhagic form, given the availability of the vector. What about *Wuchereria bancrofti*, the only filariasis to be transmitted by culicidae? Although the city of Kinshasa is located outside the area of its transmission, it is not excluded that local transmission may take place there.

Thus, as evidenced by the results of this survey, the risk of malaria transmission within the UHK itself can not be neglected. We investigated our capture in laboratory of the UHK Unit of Parasitology, and we did not find any of the mosquitoes collected were infected mainly female *Anopheles* by examining the presence of sporozoites in their salivary glands. Another study can investigate if the hospital had any patients with a mosquito-transmitted disease in the rooms where the mosquitoes were collected because it was not the aim of our study. We know by literature review that the risk of transmission of parasitic infections (yellow fever, dengue fever, Japanese encephalitis, filariasis), viral hepatitis and HIV, is very real for some patients, but that the risk of infection remains for others purely theoretical: As mosquitoes can transmit yellow fever, dengue fever and Japanese encephalitis, there is no evidence that they are capable of transmitting HIV. The level of mosquito’s nuisance found at UHK has highlighted the inadequacy of mosquito control measures in this important health facility in the DRC. Adequate mosquito control measures (drying up of stagnant water collections in and around the hospital, periodic spraying of insecticides, use of mosquito nets and other insecticide-treated materials, placement of mosquito nets in windows, etc.) should be deployed. They will not only improve the quality of sleep of patients, an important element of the quality of care, but will contribute to strengthening biosecurity measures to prevent the spread within the hospital of all germs transmitted by the mosquitoes. We are thinking particularly of the hepatitis B virus and dangerous strains of *Plasmodium falciparum* probably concentrated in the hospital.

## Conclusion

The level of mosquito’s nuisance found at UHK has highlighted the inadequacy of mosquito control measures in this important health facility in the DRC. Adequate mosquito control measures (drying up of stagnant water collections in and around the hospital, periodic spraying of insecticides, use of mosquito nets and other insecticide-treated materials, placement of mosquito nets in windows, etc.) should be deployed. They will not only improve the quality of sleep of patients, an important element of the quality of care, but will contribute to strengthening biosecurity measures to prevent the spread within the hospital of all germs transmitted by the mosquitoes. We are thinking here particularly of the hepatitis B virus and dangerous strains of *Plasmodium falciparum* probably concentrated in the hospital.

